# Exercise does not induce browning of WAT at thermoneutrality and induces an oxidative, myogenic signature in BAT

**DOI:** 10.1101/649061

**Authors:** Peter Aldiss, Jo E Lewis, Irene Lupini, David J Boocock, Amanda K Miles, Francis J P Ebling, Helen Budge, Michael E Symonds

## Abstract

**Background and aim:** Exercise training elicits diverse effects on brown (BAT) and white adipose tissue (WAT) physiology in rodents. However, these animals are typically housed below their thermoneutral zone (i.e. 28-32°C). In these conditions, BAT is chronically hyperactive and, unlike human residence, closer to thermoneutrality. Therefore, we set out to determine the effects of exercise training in obese animals at 28°C (i.e. thermoneutrality) on BAT and WAT in its basal (i.e. inactive) state.

**Methods:** Sprague-Dawley rats (n=12) were housed at thermoneutrality from 3 weeks of age and fed a high-fat diet (HFD). At 12 weeks of age half these animals were randomised to 4-weeks of exercise exercise training, i.e. swim-training (1 hour/day, 5 days per week). Metabolic assessment was undertaken during the final 48h and was followed by interscapular and perivascular BAT and inguinal (I)WAT sampling for the analysis of thermogenic genes and the proteome.

**Results:** Exercise attenuated weight gain but did not affect fat mass or general metabolic parameters (i.e. fasting insulin and glucose). Interestingly, although BAT mass was increased, there was no change in thermogenic gene expression. Differentially regulated proteins in BAT enriched gene ontology (GO) terms including 2-oxoglutarate metabolic process, cytochrome-c activity and mitochondrial respiratory chain complex IV. This was accompanied by an upregulation of multiple proteins and GO terms involved in skeletal muscle physiology suggesting an adipocyte to myocyte switch in BAT. UCP1 mRNA was undetectable in IWAT despite an increase in classical ‘browning’ markers (i.e. PGC1a and ADRB3) with exercise. Enriched GO terms in IWAT included DNA binding and positive regulation of apoptosis. Impact analysis highlighted carbon metabolism and OXPHOS pathways were regulated by exercise in BAT whilst HIF-1 signalling and cytokine-cytokine receptor interaction were among those modified in IWAT.

**Conclusion:** Exercise training reduces weight gain in obese animals at thermoneutrality and is accompanied by an oxidative, myogenic signature in BAT, rather than induction of thermogenic genes. This may represent a new, UCP1-independent pathway through which BAT regulates body weight at thermoneutrality.

## Introduction

During obesity, the accumulation of excess lipid and subsequent hypertrophy of adipocytes leads to adipose tissue (AT) dysfunction [1]. These deleterious alterations in obese AT include macrophage infiltration and apoptosis, an increase in, and secretion of, inflammatory cytokines, hypoxia and insulin resistance, all of which contribute to systemic cardiometabolic risk [1-3].

Given that sustainable weight loss is hard to achieve, improving the AT phenotype is one potential avenue to preventing the onset of diseases associated with obesity. Exercise training elicits diverse effects on both general metabolic parameters (i.e. improved insulin sensitivity) and on the AT phenotype. Following exercise training, there is a switch from a pro-inflammatory M1 to a M2 macrophage phenotype where inflammation is inhibited [4]. Increased VEGF-A and reduced AT lactate following exercise suggest an induction of AT angiogenesis and reduction in AT hypoxia whilst improving AT adipokine secretion, oxidative stress, mitochondrial biogenesis and insulin sensitivity [5-7].

More recently, there has been a focus on the role of exercise training to regulate the thermogenic programme in brown (BAT) and white AT (WAT) [8]. BAT has a high oxidative capacity similar to skeletal muscle, but utilises glucose and free fatty acids (FFA) as substrates for cold and diet-induced thermogenesis following the activation of uncoupling protein (UCP)1 [9]. Yet, how exercise training regulates BAT physiology is unclear. Exercise has been shown to induce a ‘whitening’ of BAT and reduce insulin-stimulated glucose uptake, whilst promoting the appearance of ‘beige/brite’ adipocytes in classical WAT [10-12]. This adaptation has been attributed to a range of mechanisms including the downstream actions of various myokines (e.g. meteorin-like) [13], hepatokines (e.g. fibroblast growth factor 21) [14] and metabolites (e.g. B-aminoisobutyric acid) [15]. Importantly, this occurs regardless of exercise modality (i.e. treadmill, swim training and voluntary wheel running) [16].

An important caveat, however, is that rodents subjected to exercise are typically housed at c.20°C, a temperature well below their thermoneutral zone. This impacts on a number of physiological processes including adapative thermogenesis, cardiovascular function and immune cell metabolism [17, 18]. In particular, it is an important consideration when studying BAT, which is chronically active at 20°C with UCP1+ adipocytes readily seen in WAT [19]. Therefore, we used thermoneutrality (i.e. 28°C) to closer mimic human physiology and study AT in the basal state (i.e. when UCP1 is inactive). We analysed the effects of exercise training on animals kept at thermoneutrality on both interscapular (BAT) and perivascular (PVAT) BAT having previously shown these depots to exhibit a divergent response to brief nutrient excess [20] as well as inguinal IWAT. Finally, we sought to identify how the AT proteome responds to exercise training to better understand the molecular adaptations of BAT to training at thermoneutrality.

## Methods

### Animals and exercise protocol

All studies were approved by the University of Nottingham Animal Welfare and Ethical Review Board, and were carried out in accordance with the UK Animals (Scientific Procedures) Act of 1986. Twelve male Sprague-Dawley rats aged 3 weeks were obtained from Charles River (Kent, UK) and housed immediately at thermoneutrality (c.28°C) under a 12:12-hour reverse light-dark cycle (lights off at 08:00, on at 20:00) so as to closer mimic human physiology [19], minimise animal stress and maximise data quality and translatability [21]. Animals were fed a 45% high-fat diet (824018 SDS, Kent, UK) with half of these animals randomised (http://www.graphpad.com/quickcalcs/randomize1.cfm) to 4 weeks of exercise training (Ex) at 12 weeks of age. Body weight, food and water intake (ad-lib) were monitored weekly. At 12 weeks of age, the EX group were acclimatised to water (c.35°C) for a 3-day period (10-20 minutes per day). After acclimatisation, the EX group underwent the 4-week swim training programme (1 h/day for 5 days/week).

### Metabolic cages

Animals were individually placed in an open-circuit calorimeter (CLAMS: Columbus Instruments, Linton Instrumentation, UK) for 48 h towards the end of the study. Assessment of whole body metabolism was performed as previously described [20], after which all animals were weighed and fasted overnight prior to euthanasia by rising CO_2_ gradient. BAT, IWAT and PVAT were then rapidly dissected, weighed, snap-frozen in liquid nitrogen and stored at −80°C for subsequent analysis.

### Gene expression analysis

Total RNA was extracted from each fat depot using the RNeasy Plus Micro extraction kit (Qiagen, West Sussex, UK) following an adapted version of the single step acidified phenol-chloroform method. RT-qPCR was carried out as previously described [20] using rat-specific oligonucleotide primers (Sigma) or FAM-MGB Taqman probes (see Supp Table 1 for primer list). Gene expression was determined using the GeNorm algorithm against two selected reference genes; *RPL19:RPL13a in BAT* and IWAT (stability value M = 0.26 in BAT and 0.224 in IWAT) and RPL19:HPRT1 in PVAT (stability value M = 0.209).

**Table 1.**
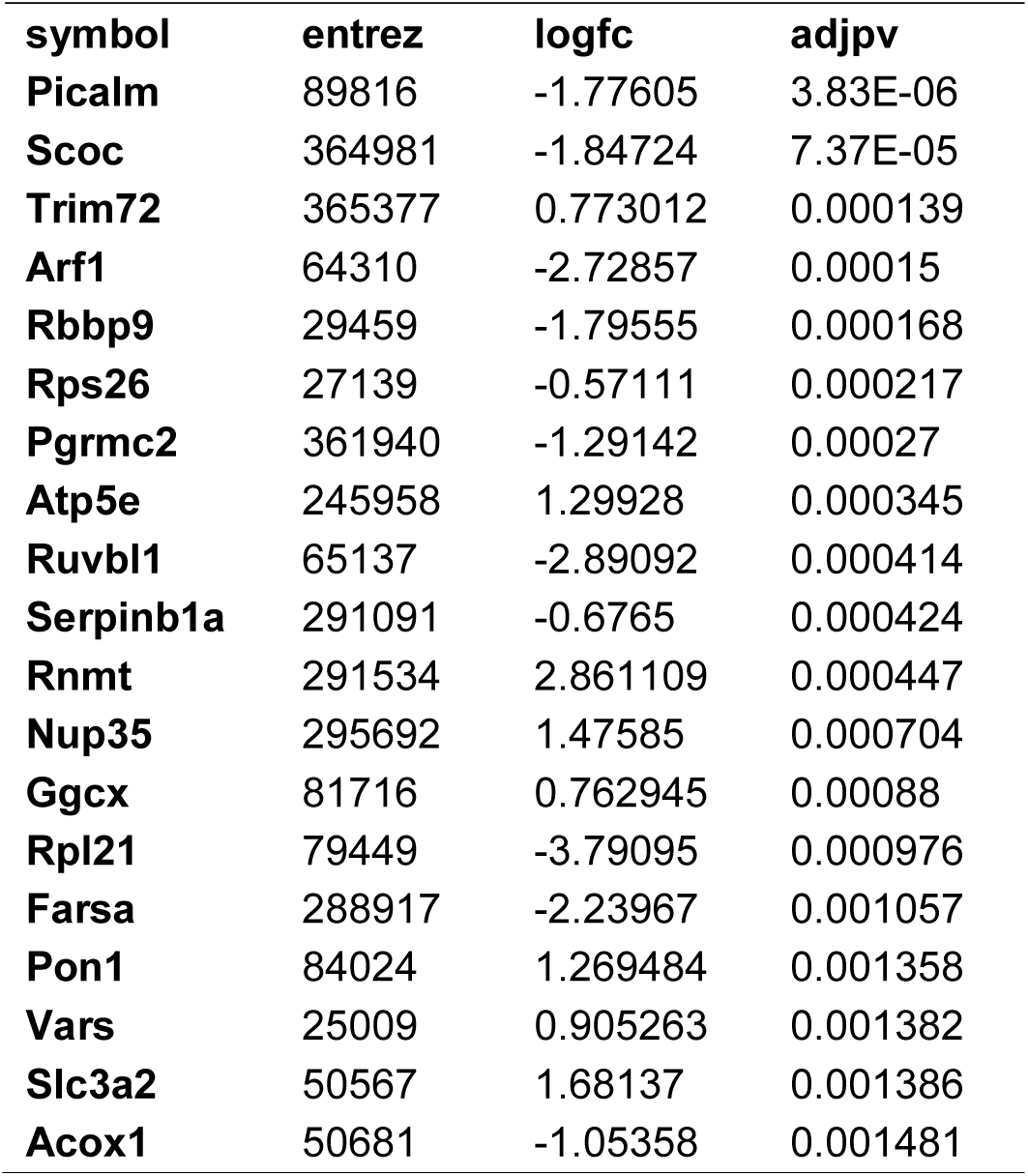
Top 20 differentially regulated proteins in BAT

### Serum analysis

Serum was thawed gently on ice with concentrations of glucose (GAGO-20, Sigma Aldrich, Gillingham, UK), triglycerides (LabAssay ™ Trigylceride, Wako, Neuss, Germany), non-esterified fatty acids (NEFA-HR(2), Wako, Neuss, Germany), insulin (80-INSRT-E01, Alpco, Salem, NH, USA) and leptin (EZRL-83K, Merck, Darmstadt, Germany) were measured following manufacturer’s instructions.

### Mass spectrometry

Protein extraction, clean up and trypsinisation was carried out as previously described [20]. Briefly, 50-100 mg of frozen tissue was homogenised in 500 μL CellLytic MT cell lysis buffer (Sigma, C3228) prior to removal of lipid and other contaminants using the ReadyPrep 2D cleanup Kit (Biorad, 1632130). Samples were then subjected to reduction, alkylation and overnight trypsinisation, following which they were dried down at 60°C for 4 h and stored at 80°C before resuspension in LCMS grade 5% acetonitrile in 0.1% formic acid for subsequent analysis. Analysis by mass spectrometry was performed on a SCIEX TripleTOF 6600 instrument as previously described [22]. Briefly, samples were analysed in both SWATH (Data Independent Acquisition) and IDA (Information Dependent Acquisition) modes for quantitation and spectral library generation respectively. IDA data was searched together using ProteinPilot 5.0.2 to generate a spectral library and SWATH data was analysed using Sciex OneOmics software [23] extracted against the locally generated library as described previously [20].

### 2.6 Statistical analyses

Statistical analyses were performed in GraphPad Prism version 8.0 (GraphPad Software, San Diego, CA). Data are expressed as Mean ± SEM with details of specific statistical tests in figure legends.

Functional analysis of the proteome (fold change ± 0.5 and OneOmics confidence score cut-off of 0.75) was performed using the Advaita Bioinformatic iPathwayGuide software (www.advaitabio.com/ipathwayguide.html). Significantly-impacted biological processes, molecular interactions and pathways were analysed in the context of pathways obtained from the Kyoto Encyclopedia of Genes and Genomes (KEGG) database (Release 84.0+/10-26, Oct 17) [24] and the Gene Ontology Consortium database (2017-Nov) [25]. The Elim pruning method, which removes genes mapped to a significant GO term from more general (higher level) GO terms, was used to overcome the limitation of errors introduced by considering genes multiple times [26]. Analysis of protein-protein interactions (PPI) and GO term enrichment of these PPI networks was performed using NetworkAnalyst (www.networkanalyst.ca).

## Results

### Exercise-training increases BAT mass and regulates thermogenic genes in perivascular and inguinal WAT

Swim training in diet-induced obesity did not affect body weight or total fat mass but significantly attenuated weight gain during the 4-week intervention period (78.6 ± 5.3 vs 52 ± 7.7g, p=0.03; Fig. 1A-C) without effect on serum hormones or metabolites (Fig 1. J-M), hepatic weight or hepatic triglycerides (Fig 1. N-O). Despite attenuated weight gain, BAT mass increased (Fig. 1D) although no change in key thermogenic (e.g. UCP1) or lipogenic (FASN) genes (Fig. 1P) was detected. Interestingly, UCP1 was upregulated in PVAT along with PGC1a, a marker of mitochondrial biogenesis and P2RX5, a purinergic receptor and brown/beige adipocyte cell surface marker (Fig. 1P). Similarly, genes governing fatty-acid oxidation (PPARA) and lipogenesis (FASN) were upregulated in PVAT. With regards to ‘browning’, UCP1 mRNA was undetectable in IWAT and was not induced with exercise training despite an upregulation of PGC1a, ADRB3, DIO2 and PPARA (Fig. 1P).

**Figure 1.**
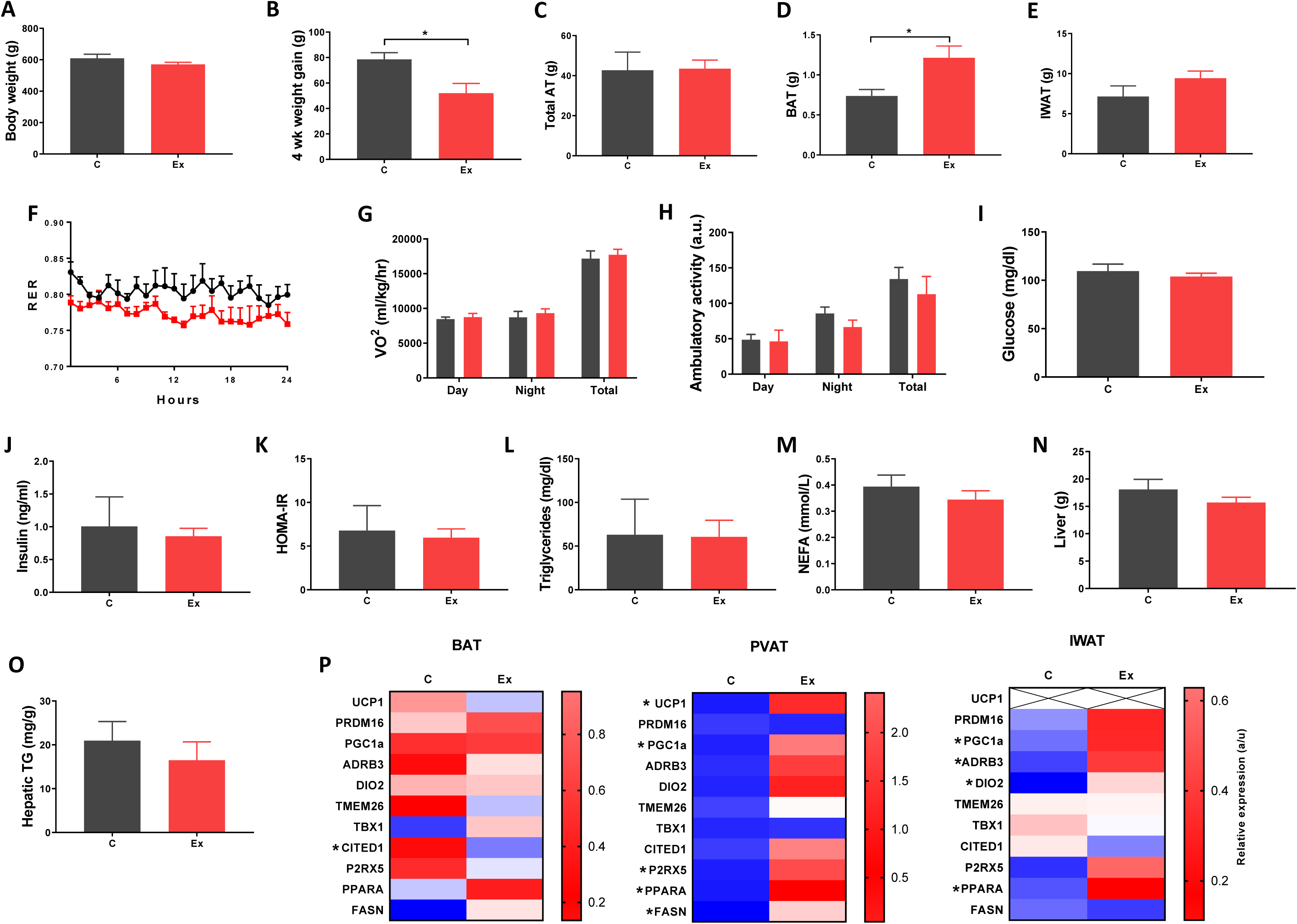
Exercise training (Ex) attenuated weight gain and increased brown adipose tissue (BAT) mass but had no effect on insulin, glucose, triglycerides or non-esterified fatty acids. (A) Final body weight, (B) 4 week intervention weight gain, (C) total fat mass, (D) BAT mass, (E) inguinal white adipose tissue (IWAT) mass, (F) respiratory exchange ratio (RER), (G) oxygen consumption (VO_2)_, (H) ambulatory activity, (I-E) serum hormones and metabolites, (N-O) liver weight and hepatic triglycerides and (P) thermogenic genes in BAT, PVAT and IWAT. Data expressed as mean ± SEM, n=4-5 per group. For comparison, data was analysed by either Students t-test (A-E and I-P) or two-way ANOVA (F-H) and Sidak post-hoc tests. Significance denoted as * <0.05; ** <0.01 or *** <0.001.

### Identification of differentially regulated proteins in BAT and IWAT in response to swim-training

We then sought to determine the exercise-induced effects on the proteome of these BAT and IWAT depots. We identified 353 differentially regulated proteins in BAT (Table 1: Top 20 proteins; Supp. Table 1: Full list). The most significantly altered proteins were involved in mitochondrial ATP synthase (ATP5E), nucleopore (NUP35), ADP ribosylation (ARF1 and SCOC) and progesterone binding (PGMRC2). Among the proteins most upregulated in BAT were those involved in assembly of the skeletal muscle cytoskeleton (PDLIM3 and MYH4), muscle contraction (TNNI2) and muscle-specific phosphoglycerate mutase metabolism (PGAM2). Proteins involved in calcium sensing in the luminal sarcoplasmic reticulum (CASQ1) and beta adrenergic signalling (CAPN1 and PSMB7) were found to be the most downregulated in BAT. Conversely, only 189 proteins were differentially regulated in IWAT after exercise. The most significantly altered proteins (Table 2: Top 20 proteins; Supp. Table 2: Full list) were involved in the trafficking of GLUT4 (TUSC5), the mitochondrial electron transport chain (NDUFS6), beta adrenergic signalling (PSMB7), TLR4 signalling (LRRFIP2) and apoptosis (ATG7 and BIN1). Proteins with the greatest fold change were those involved in cell adhesion (CD44 and VCAN), FFA and lipoprotein metabolism (FABP3 and APOC1), TGF-beta signalling (TSC22D1), purine and mitochondrial metabolism (LHPP and COX6A1) and the acetylation of nucleosomes and DNA binding (HET and HMGB1).

**Table 2.**
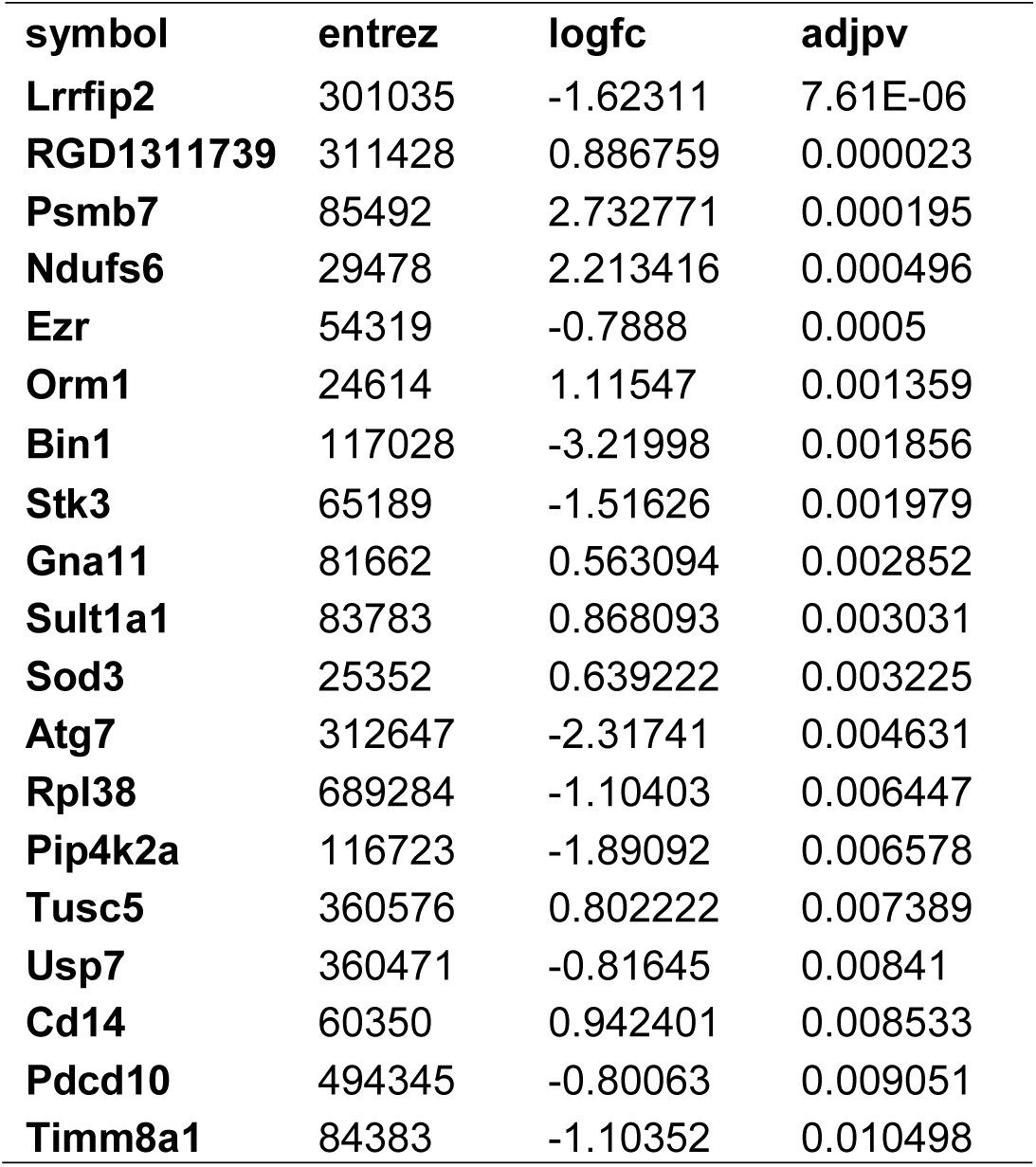
Top 20 differentially regulated proteins in IWAT

### Exercise training enriches mitochondrial and skeletal muscle related GO terms and pathways in BAT

We then carried out functional analysis of the BAT and IWAT proteome. The differentially regulated proteins in BAT enriched GO terms (Table 3; Supp. Table 3: Full list) including *2-oxoglutarate metabolic process, generation of precursor metabolites, cytochrome-c oxidase activity, mitochondrial respiratory chain complex IV* and *proton transporting ATP synthase activity* (Fig. 2A-D). There was also an enrichment of GO terms related to skeletal muscle physiology including *sarcomere, myosin complex* and *skeletal muscle tissue development* (Fig. 2E-G). In IWAT, (Table 4; Supp. Table 4 for full list) the differentially regulated proteins enriched GO terms including *positive regulation of apoptotic process, positive regulation of ATPase activity* and *lipid droplet* (Fig. 3A-C). In addition, a number of GO terms associated with RNA processing were enriched including *spliceosomal complex* and *negative regulation of transcription from RNA polymerase II promoter* (Fig. 3D-E).

**Table 3.**
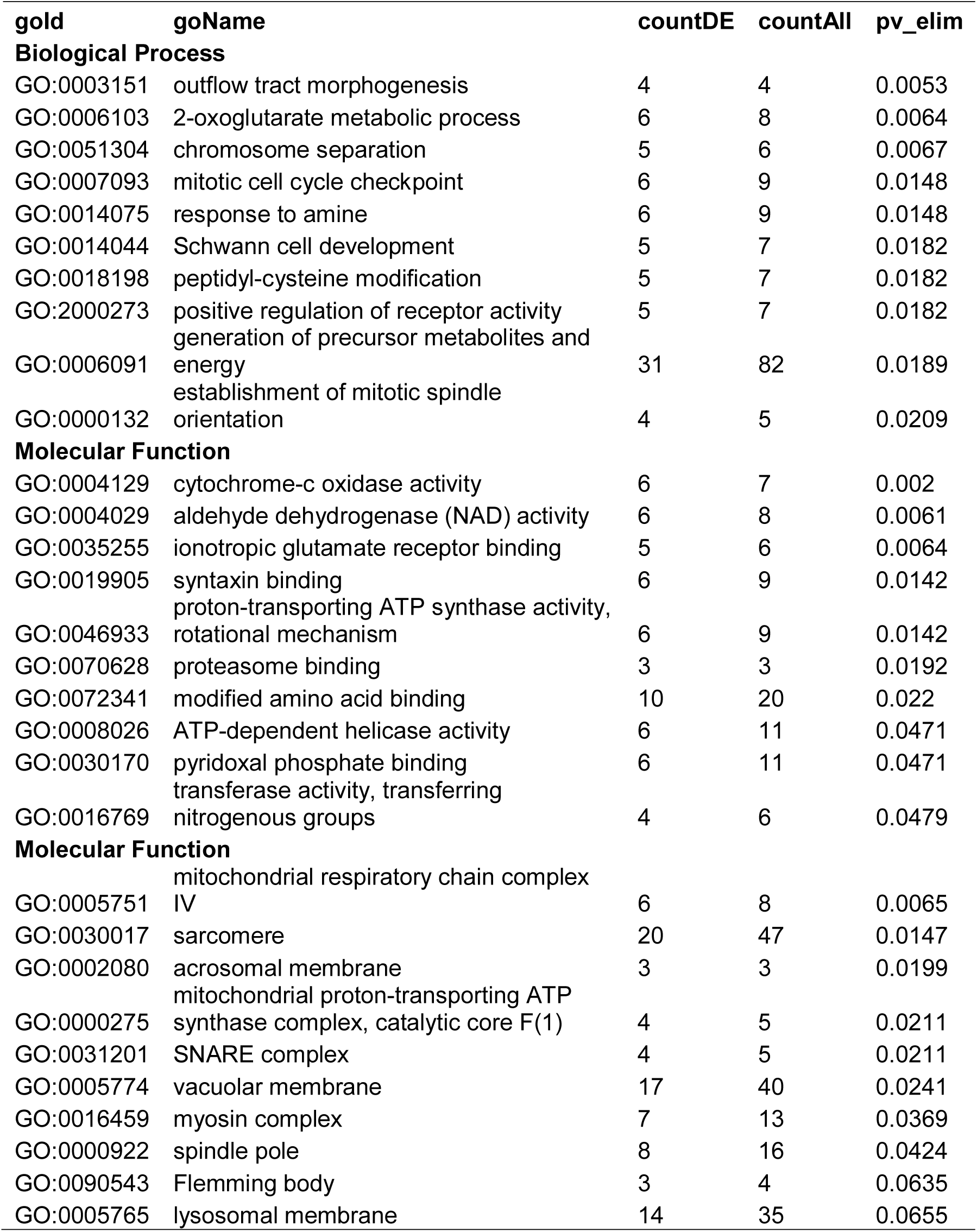
GO terms enriched in BAT.

**Table 4.**
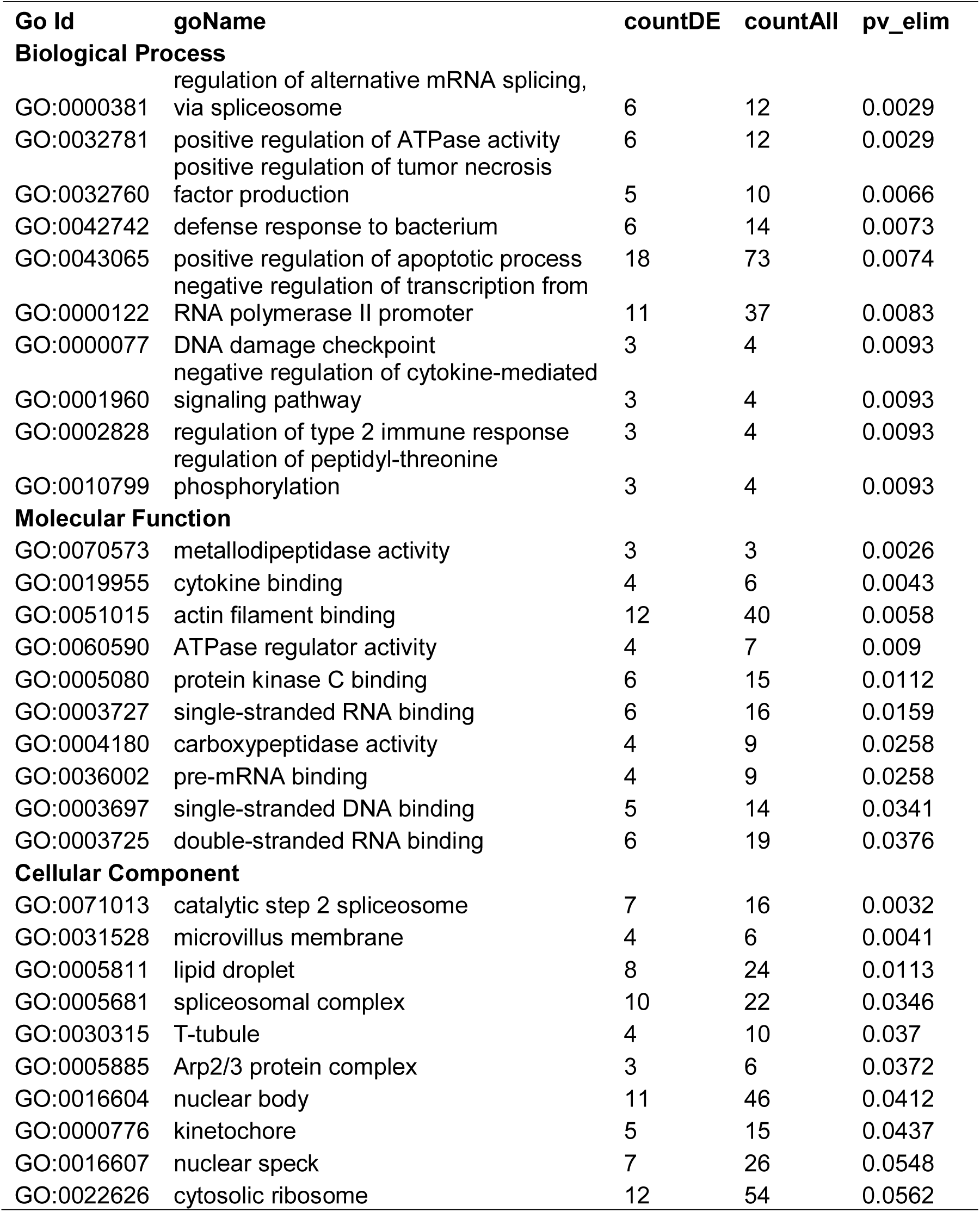
GO terms enriched in IWAT.

**Figure 2.**
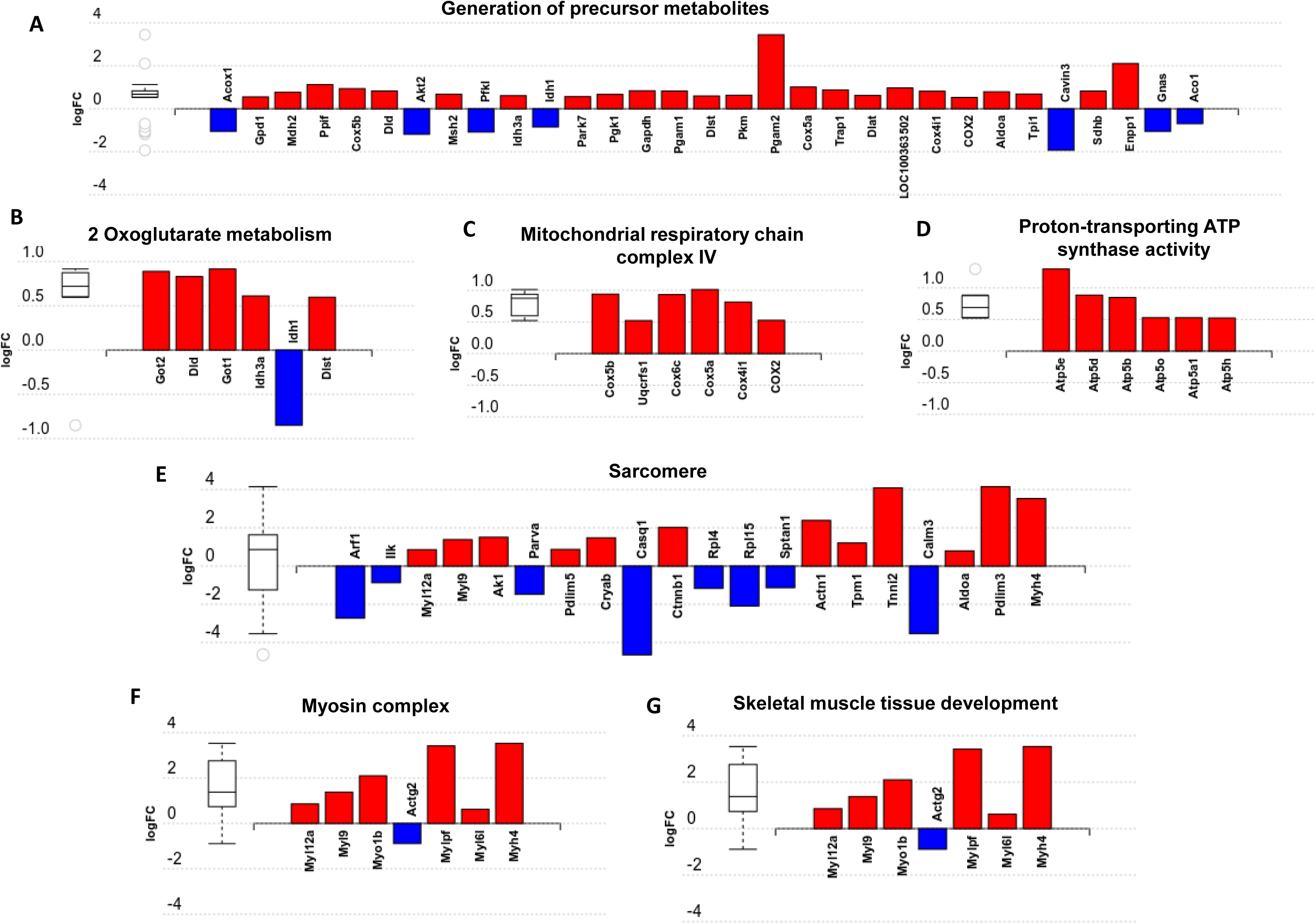
Overview of enriched metabolic and skeletal muscle related gene ontology (GO) terms in brown adipose tissue. (A) generation of precursor metabolites, (B) 2 Oxoglutarate metabolism, (C) Mitochondrial respiratory chain complex IV, (D) Proton-transporting ATP synthase activity, (E) Sarcomere, (F) Myosin complex and (G) Skeletal muscle tissue development. Figures created with AdvaitaBio IPathwayGuide.

**Figure 3.**
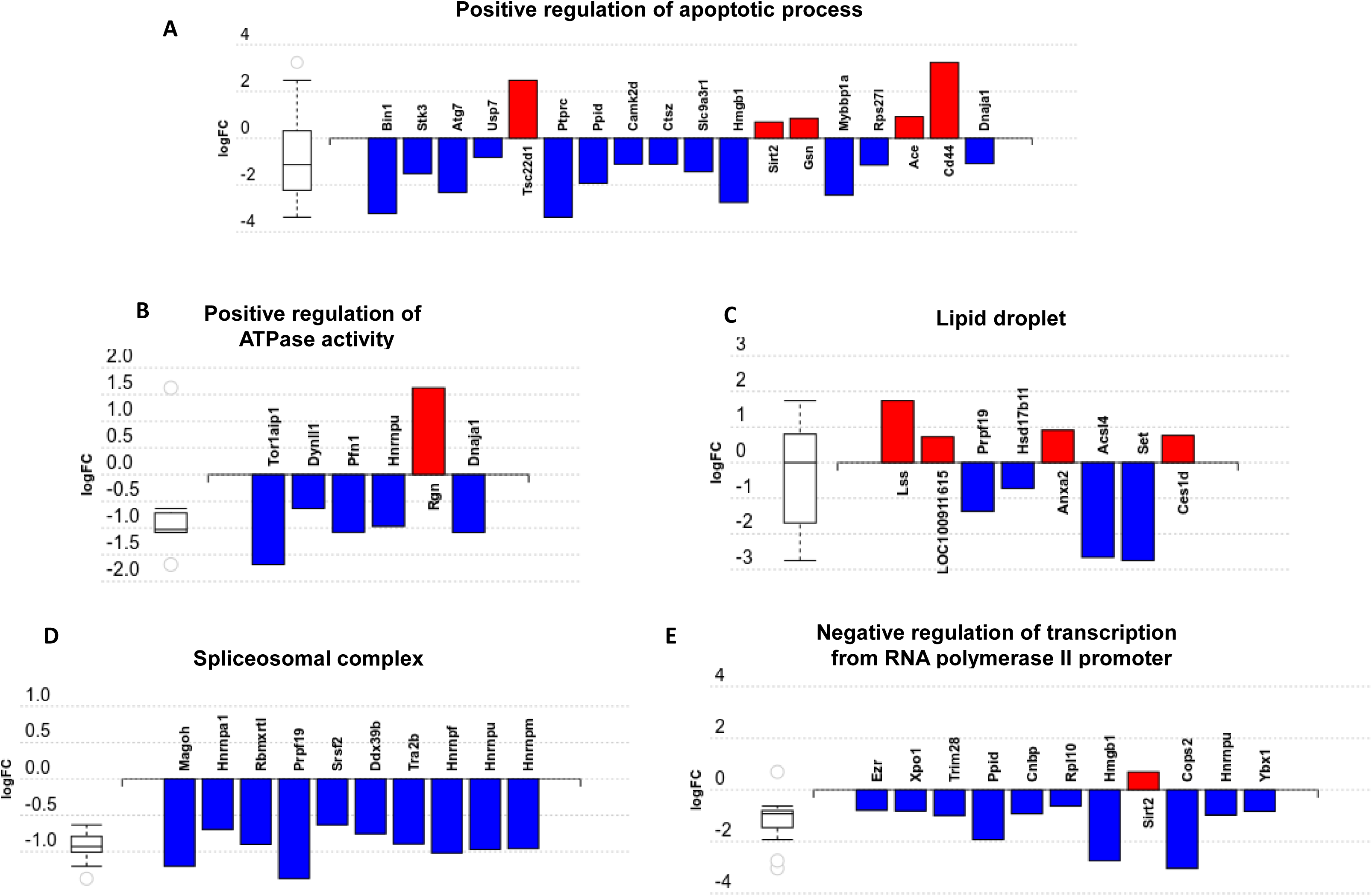
Overview of enriched gene ontology (GO) terms in inguinal white adipose tissue. (A) Positive regulation of apoptotic process, (B) Positive regulation of ATPase activity, (C) Lipid droplet, (D) Spliceosomal complex and (E) Negative regulation of transcription from RNA polymerase II promoter. Figures created with AdvaitaBio IPathwayGuide.

Impact analysis, which combines both classical overrepresentation analysis with the perturbation of a pathway, highlighted several metabolic pathways modified by exercise (Supp. Table 5) including *ascorbate and aldarate metabolism, biosynthesis of amino acids, glycolysis/gluconeogenesis* and *carbon metabolism*. Lipid related pathways such as *glycerophospholipid metabolism* and the *sphingolipid signalling pathway* were also impacted alongside those associated with mitochondrial proteins such as *Alzheimer’s disease* and *oxidative phosphorylation* (Fig. 4A, C and E). In IWAT, the impacted pathways (Fig. 4B and D) included the *spliceosome and Fc gamma R-mediated phagocytosis*.

**Figure 4.**
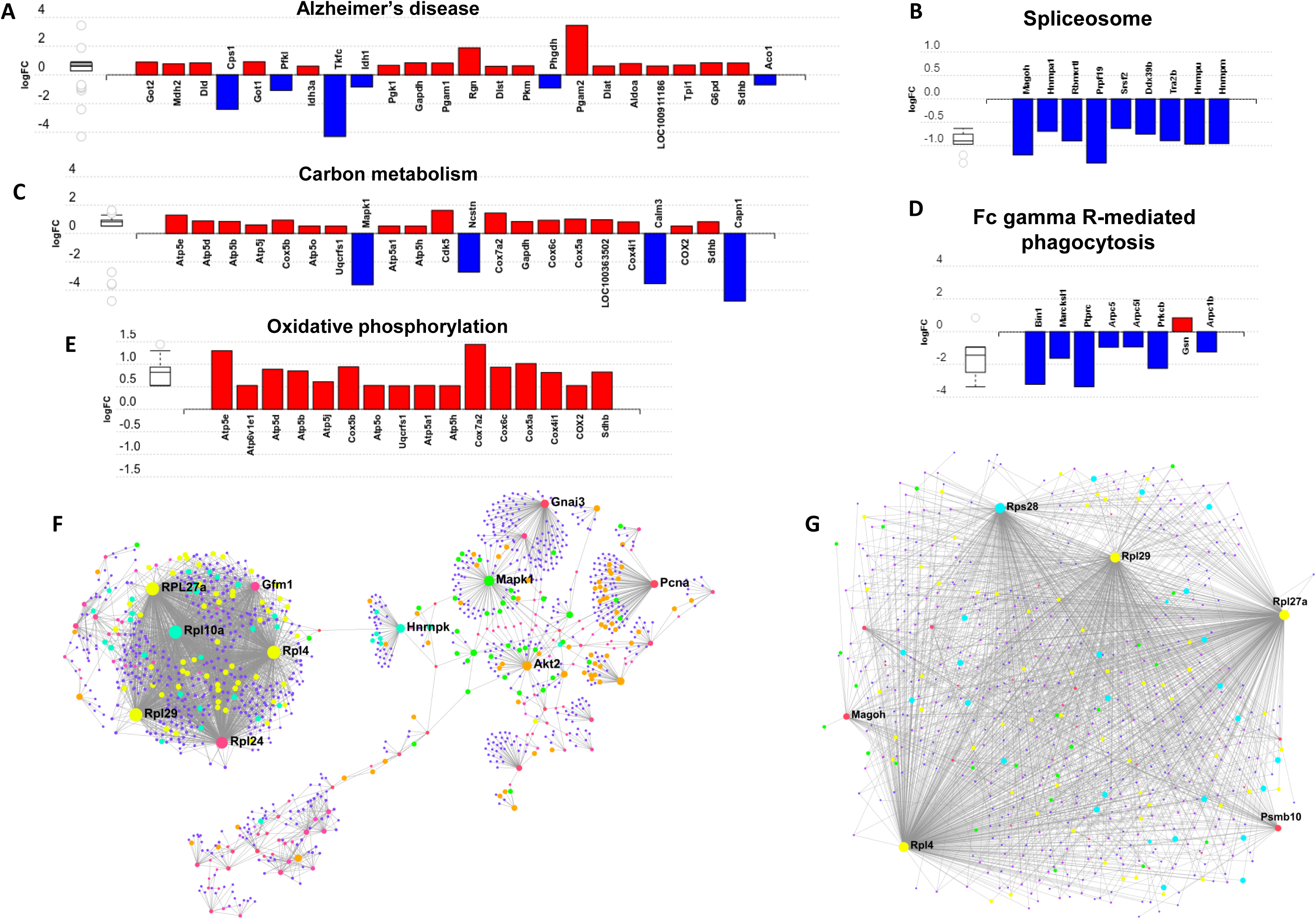
Overview of significantly impacted pathways and protein-protein interaction networks in brown adipose tissue (A, C, E and F) and inguinal white adipose tissue (B, D and G). (A) Alzheimer’s disease, (B) Spliceosome, (C) Carbon metabolism, (D) Fc gamma R-mediated phagocytosis and (E) Oxidative phosphorylation. (F) Protein-protein interaction in BAT and (G) IWAT. Figures A-E created with AdvaitaBio IPathwayGuide. Figures F-G created with NetworkAnalyst.

### Characterisation of the ‘interactome’ in exercise-trained brown and white adipose tissues

To better understand how our differentially altered proteins affect downstream signalling pathways, we characterised the ‘interactome’ of BAT and IWAT through analysis of protein-protein interactions. NetworkAnalyst generated 28 sub-networks (i.e. the main ‘continent’ and 27 ‘islands’) in BAT with the main network consisting of 1091 proteins. Hub proteins in this main network included the ribosomal proteins (i.e. RPL27a and RPL10a), the mitochondrial elongation factor GFM1, AKT Serine/Threonine Kinase 2 (Akt2) and mitogen-activated protein kinase 1 (MAPK1). Further analysis demonstrated interacting proteins in this main network enriched 30 biological processes (Supp. data PPI, Table 2) including *chromatin assembly or disassembly, developmental growth and muscle organ development* and 30 molecular functions (Fig. 4F) including *RNA binding* (yellow), *steroid dehydrogenase activity* (green), *neuropeptide hormone activity* (orange) and *transcription cofactor activity* (light blue).

In IWAT, the ‘interactome’ was smaller, consisting of 19 sub-networks (i.e. 1 ‘continent’ and 18 ‘islands’) with the main network made up of 488 proteins. Hub proteins in the main network again included multiple ribosomal proteins (i.e. RPL27a and RPL4) in addition to Proteasome subunit B10 (PSMB10) and the spliceosomal protein mago homolog, exon junction complex subunit (MAGOH). Further analysis demonstrated that the interacting proteins in this main network were involved in 6 biological processes (Supp. data PPI, Supp. Table 6) including *chromatin assembly or disassembly, sensory taste perception* and *RAS protein signal transduction* and 5 molecular functions (Fig. 4G) including *RNA binding (yellow), transcription cofactor activity (light blue)* and nucleotide *binding (green)*.

## Discussion

The vast majority of studies investigating the function of BAT in rodents have been carried out at temperatures (i.e. c.20-22°C) which are well below thermoneutrality (i.e. c.28-33°C). These environmental differences have diverse effects on physiology, immunity and metabolism [17, 18]. Whilst the use of thermoneutrality has been suggested as the optimal environment to mimic human physiology there is ongoing debate as to ‘how high’ we should go [18, 27, 28]. Here, using thermoneutral housing, we show exercise training induces an oxidative phenotype in BAT of obese animals that is associated with an enrichment of GO terms involved in skeletal muscle physiology and of multiple pathways associated with altered mitochondrial metabolism (i.e. Alzheimer’s disease and oxidative phosphorylation). Unlike studies conducted at sub-thermoneutrality [16, 29], we show UCP1 mRNA is absent in IWAT of obese animals raised at thermoneutrality and is not induced with exercise training. Instead, IWAT in exercise trained animals exhibits a reduction in apoptotic proteins and perturbations in the spliceosomal pathway.

### A thermogenic response in PVAT and adipocyte-to-myocyte switch in BAT

Despite no induction of thermogenic genes in BAT, an upregulation of these markers in PVAT suggests an uncoupling of the response to exercise training in anatomically distinct BAT depots. A downregulation of thermogenic genes and ‘whitening’ of thermogenic AT has previously been attributed to increases in core body temperature with exercise training [8]. This would seem not to be the case, however, given an increase of these genes in PVAT which plays a critical role in heating blood prior to circulation [30]. The physiological role of these alterations in PVAT is unclear but our data suggests that exercise training induces a thermogenic response in PVAT, whilst BAT shifts towards a skeletal muscle phenotype.

A myogenic signature in brown adipocytes was first established in 2007 when it was shown that the expression of myogenic genes in differentiating brown adipocytes was a characteristic that clearly distinguished them from white adipocytes [31]. It was subsequently shown that both BAT and skeletal muscle derive from the same Pax7+ / Myf5+ progenitor cells and that the transcription factor PRDM16 drives the fate of these progenitors to committed brown adipocytes [32]. Despite these shared characteristics, a definitive physiological role for these myogenic proteins in BAT has not been demonstrated. In WAT, blockade of the β3-receptors induces myogenesis with a) the emergence of MyoD+ mononucleated cells which undergo myogenesis even under adipogenic conditions and b) the fusion of stromal cells which form multinucleated myotubes that ‘twitch’ and express myosin-heavy chain (MHC) [33]. A subset of UCP1+ beige adipocytes (c.15% of total beige adipocytes) in Myod1-CreERT2 reporter mice are derived from MyoD+ cells located adjacent to the microvasculature, and this ‘glycolytic’ beige fat exhibits enhanced glucose metabolism compared to typical beige adipocytes.

Given that exercise training drives myogenesis in skeletal muscle, it may have a similar impact on BAT given their shared developmental origins. The enrichment of pathways involved in amino acid metabolism (i.e. *biosynthesis of amino acids*; *glycine, serine, threonine, arginine and proline metabolism*) seen in the current study would certainly point towards this. Mice lacking interferon regulatory factor 4 (IRF4) in BAT (BATI4KO) exhibit reduced exercise capacity at both low and high-intensity treadmill running, and display selective myopathy [34]. Interscapular BAT of these exercise intolerant mice is characterised by an upregulation of genes governing skeletal muscle physiology, including MyoD1, troponin T1 and myostatin. This suggests that myogenesis in BAT may have an adverse effect on whole body physiology. Alongside the induction of myogenic proteins was an upregulation of proteins involved in the *generation of precursor metabolites* and *energy and mitochondrial respiration*. Phosphoglycerate mutase (PGAM) is a key glycolytic enzyme, and mutations in the muscle isoform of this gene (i.e. muscle phosphoglycerate deficiency) cause tubular aggregates in muscle and exercise intolerance. An upregulation of the muscle isoform of this protein (PGAM2) in BAT further strengthens the idea of a switch towards a muscle phenotype and, along with the upregulation in mitochondrial proteins, points towards increased glycolytic capacity. An enrichment of multiple mitochondria associated metabolic pathways including OXPHOS further suggests that, despite no demonstrable impact on UCP1 in BAT, the metabolic activity of this tissue has, nevertheless, increased. Whilst further work is needed to validate and corroborate this data, we propose this as a novel, UCP1-independent pathway through which exercise regulates BAT metabolism.

### A role for exercise in the attenuation of adipose tissue apoptosis

Whilst prior work has demonstrated an induction of thermogenic genes in WAT following exercise training, we show that UCP1 mRNA is absent in IWAT of animals raised at thermoneutrality from weaning. Instead, there are alterations to apoptotic and spliceosomal proteins. Dysregulated apoptotic processes are associated with AT inflammation and insulin resistance [35, 36]. Here, we show one potential benefit of exercise training on IWAT is a downregulation of multiple proteins governing the ‘*positive regulation of apoptotic process*’. Bridging integrator (BIN)1, for instance, is a MYC proto-oncogene interacting factor that activates caspase-independent apoptosis in cancer cells, though its role in AT immunometabolism is unknown [37]. Autophagy-related (ATG)7 is a ubiquitin activating enzyme which forms a complex with caspase-9 to cross-regulate autophagy and apoptosis [38]. Adipocyte specific ATG7 k/o mice are lean with reduced fat mass, increased insulin sensitivity, an increase in BAT thermogenesis and are resistant to diet-induced obesity [39, 40]. Other proteins of particular interest include MYB binding protein (MYBBP)1a and CD44. MYBBP1a is a SIRT7 interacting protein which regulates nucleolar stress and ribosomal DNA synthesis that is increased in visceral AT of obese mice and negatively regulates adipogenesis [41, 42]. Downregulation of MYBBP1a by exercise training may be a putative mechanism whereby physical activity and/or exercise regulates adipocyte number and size. Finally, CD44 mRNA is three-fold higher in AT of insulin-resistant humans and correlates with CD68 and IL6, whilst CD44 k/o mice are phenotypically healthier and exhibit reduced AT inflammation [43, 44].

Impact analysis demonstrated a number of significantly perturbed pathways. Epidermal growth factor (EGF) was the single protein differentially regulated in pathways including melanoma, phospholipase D signalling and PI3K-Akt signalling pathways. The role of EGF in adipogenesis is well described with EGF receptor (ErbB1) abundance reduced in insulin resistant women with Type 2 diabetes. Importantly, EGF exerts insulin-like effects on adipocytes and skeletal muscle and this exercise-induced increase may be one way in which physical exercise potentiates insulin-sensitising effects in these tissues [45]. The most impacted pathway, however, was the spliceosome. This multi-megadalton ribonucleoprotein complex removes introns from RNA polymerase II transcripts (pre mRNAs) and is a crucial step in mRNA synthesis [46]. Given that c.95% of genes are subject to alternative splicing, a downregulation of proteins involved in the spliceosome pathway would likely have major downstream effects on AT function, and may be driving the alterations observed in the proteome [47]. Peturbation of the spliceosomal pathway was associated with an enrichment of the GO term ‘*regulation of RNA metabolic process*’ in which 36 of 44 proteins were reduced. Why exercise training downregulates large numbers of proteins involved in pre-mRNA synthesis and in RNA metabolism is, however, unclear and merits further investigation.

### Characterising the exercise interactome in BAT and WAT

Analysis of protein-protein interactions is an important step in understanding the communication between proteins and identification of putative signalling pathways occurring in specific tissues following an intervention. Here, we have identified multiple ribosomal proteins (i.e. RPL27a and RPL10a), AKT2 and MAPK1 as important hub proteins controlling the regulation of chromatin assembly, muscle organ development and steroid hormone dehydrogenase activity in exercised BAT. It has been suggested that increased ribosomal biogenesis plays a major role in skeletal muscle hypertrophy and we propose that these specific ribosomal hub proteins play a role in the shift towards a myogenic signature in BAT [48]. Ribosome biogenesis is regulated by growth factors, and IGF1 specifically regulates muscle hypertrophy through both the PI3K-AKT-mTOR and MAPK signalling pathways, with the latter playing a key role in regulating ribosomal biogenesis [49, 50]. Furthermore, PCNA, is a cell cycle protein, whose upregulation suggests satellite cells have entered the cell cycle [51, 52]. That AKT2, MAPK1 and PCNA are identified as hub proteins suggests that both them, and their interacting proteins, are important to the induction of myogenic proteins in BAT. These proteins and their downstream targets offer novel insight into the induction of myogenesis in BAT and the regulation of BAT physiology with exercise. The identification of ribosomal proteins as both hub and interacting proteins in WAT suggests they are also important in the general regulation of adipose tissue physiology by exercise training, though their role in WAT is less clear. To date, no functional role for these hub proteins in AT has been shown though the identification of MAGOH as a hub protein suggests it may play a key role in the regulation of the spliceosome pathway. Functional studies of these hub proteins and signalling pathways in AT is needed to better understand how exercise regulates WAT biology.

## Conclusion

We propose that BAT is dormant at thermoneutrality and, with exercise training, undergoes an adipocyte-to-myocyte switch towards a muscle phenotype. The physiological relevance of this adaptation is unclear and BAT-muscle crosstalk, in particular, merits further investigation. Meanwhile, WAT exhibits a reduction in apoptotic proteins and a wholesale downregulation of proteins involved in pre-mRNA synthesis and RNA metabolism.

## Supporting information

Supp data PPI

## Acknowledgements

The study was funded by he British Heart Foundation [grant number FS/15/4/31184/].

